# Qualitative and quantitative top-down proteomics of human colorectal cancer cell lines identified 23000 proteoforms and revealed drastic proteoform-level differences between metastatic and non-metastatic cancer cells

**DOI:** 10.1101/2021.10.27.466093

**Authors:** Elijah N. McCool, Tian Xu, Wenrong Chen, Nicole C. Beller, Scott M. Nolan, Amanda B. Hummon, Xiaowen Liu, Liangliang Sun

## Abstract

Understanding cancer metastasis at the proteoform level is crucial for discovering new protein biomarkers for cancer diagnosis and drug development. Proteins are the primary effectors of function in biology and proteoforms from the same gene can have drastically different biological functions. Here, we present the first qualitative and quantitative top-down proteomics (TDP) study of a pair of isogenic human metastatic and non-metastatic colorectal cancer (CRC) cell lines (SW480 and SW620). This study pursues a global view of human CRC proteome before and after metastasis in a proteoform specific manner. We identified 23,319 proteoforms of 2,297 genes from the CRC cell lines using capillary zone electrophoresis-tandem mass spectrometry (CZE-MS/MS), representing nearly one order of magnitude improvement in the number of proteoform identifications from human cell lines compared to literature data. We identified 111 proteoforms containing single amino acid variants (SAAVs) using a proteogenomic approach and revealed drastic differences between the metastatic and non-metastatic cell lines regarding SAAVs profiles. Quantitative TDP analysis unveiled statistically significant differences in proteoform abundance between the SW480 and SW620 cell lines on a proteome scale for the first time. Ingenuity Pathway Analysis (IPA) disclosed that many differentially expressed genes at the proteoform level had diversified functions and were closely related to cancer. Our study represents a milestone in TDP towards the definition of human proteome in a proteoform specific manner, which will transform basic and translational biomedical research.

**For TOC only:** 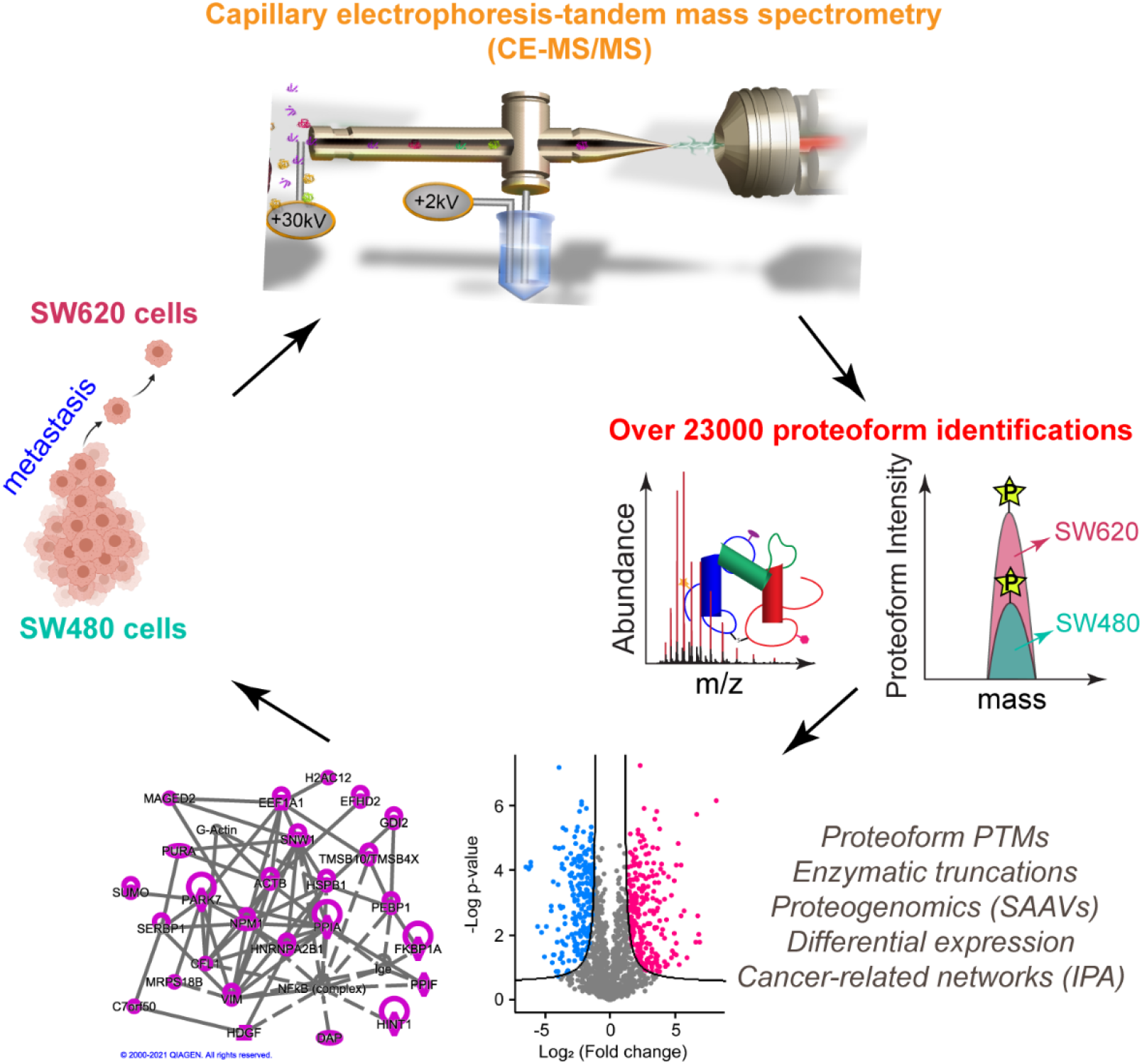

---

Colorectal cancer (CRC) is the third most common cancer worldwide and has a high mortality rate even with recent improvements in therapies.^1,2^ CRC metastasis is the main cause of CRC-related death. New insights into the molecular mechanisms of CRC metastasis will undoubtedly be beneficial for developing more effective drugs.^3–5^ Extensive studies have been completed with the goal of understanding CRC metastasis at the transcriptome level, generating tremendous information about the landscape of mRNA across different stages of CRC. ^[6,7]^ However, nucleic-acid–based measurements do not correlate with protein abundance, which are the primary effectors of function in biology.^[8]^ Quantitative bottom-up proteomics (BUP) studies of metastatic and non-metastatic CRC cell lines have been completed to discover new protein regulators involved in CRC metastasis.^[4,9,10]^ Unfortunately, those BUP studies provided limited information of the proteoforms. These proteoforms represent all possible protein molecules derived from the same gene resulting from genetic variations, RNA alternative splicing, and protein post-translational modifications (PTMs).^[11]^ It has been well demonstrated that proteoforms from the same gene can have drastically different biological functions.^[12]^ Mass spectrometry (MS)-based top-down proteomics (TDP) directly measures intact proteoforms and delineates proteomes in a proteoform-specific manner.^[13]^ TDP is invaluable for pursuing a better understanding of molecular mechanisms of cancers and discovering new proteoform biomarkers for more reliable diagnosis and drug development. ^[14]^

Here, we performed the first deep TDP study of metastatic (SW620) and non-metastatic (SW480) human CRC cell lines, aiming to produce a comprehensive proteoform-level view of the two isogenic CRC cell lines and discover novel proteoform biomarkers of CRC metastasis. We fractionated SW480 and SW620 cell lysates using one-dimensional liquid chromatography (1D-LC, size exclusion chromatography (SEC) or reversed-phase LC (RPLC)) and 2-D LC (SEC-RPLC), followed by capillary zone electrophoresis (CZE)-tandem MS (MS/MS) analyses of all the LC fractions from both cell lines for proteoform identification (ID) and label-free quantification (LFQ), **Figure 1**. The TopPIC (version 1.4.0) software was used for data analysis.^[15]^ The experimental details are described in **Supporting Information I**.

**Figure 1.**
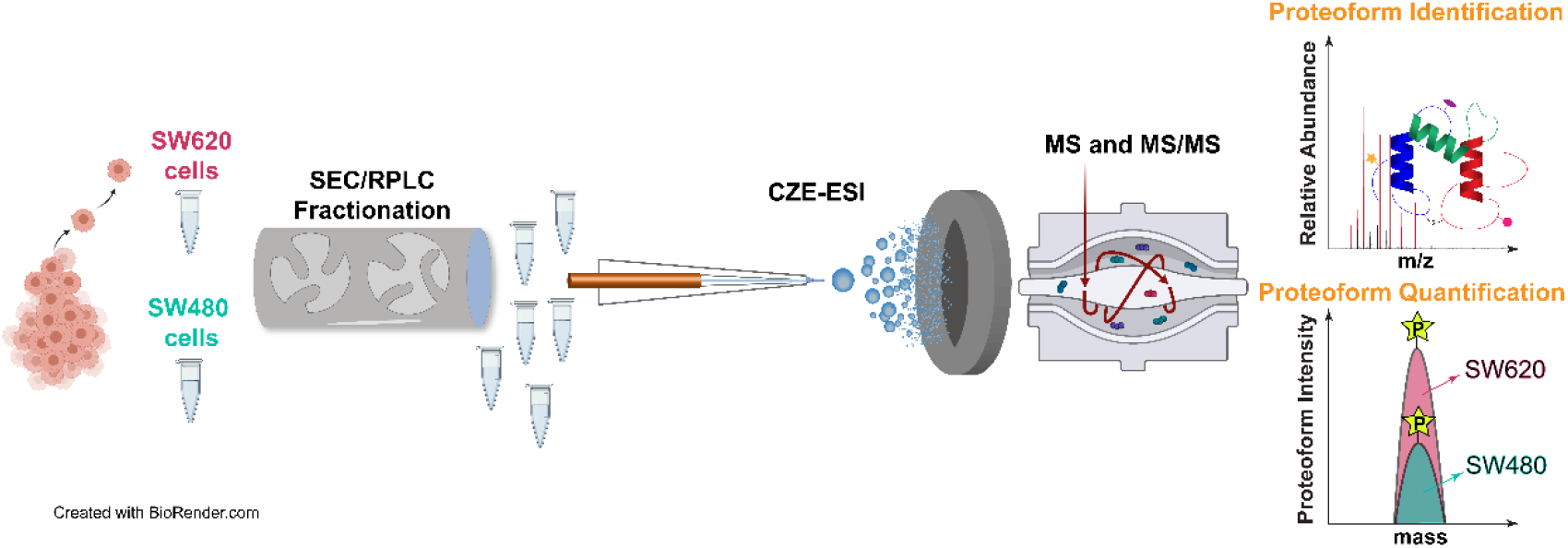
Schematic design of the TDP study of metastatic (SW620) and non-metastatic (SW480) CRC cells using LC fractionation and CZE-ESI-MS and MS/MS for proteoform identification and quantification.

One long-term goal of TDP is to characterize all the millions of proteoforms in the human body.^[16,17]^ During the last decade, because of the improvement of proteoform sample preparation, LC and CZE separation, MS and MS/MS, about 3,000 proteoforms corresponding to roughly 1,000 genes can be identified in one TDP study from human cell lines using LC-MS/MS-based platforms,^[18–20]^ and up to 6,000 proteoform IDs corresponding to 850 genes have been reported from an *E. coli* sample using a CZE-MS/MS-based workflow.^[21]^ In this study, we collected 410 MS raw files and identified in total 23,319 proteoforms derived from 2,297 genes from the SW480 and SW620 human CRC cell lines with a 1% proteoform-level false discovery rate (FDR), representing nearly one order of magnitude improvement in the number of proteoform IDs from human cell lines, **Figures 2A. Figure 2B** shows a learning curve of the number of proteoform IDs from complex proteomes using TDP by comparing this study with previous works.^[18–22]^ The data clearly demonstrate the power of our CZE-MS/MS-based TDP workflow for comprehensive characterization of proteoforms in complex proteome samples. We attribute the drastic improvement of proteoform IDs to the high separation efficiency of CZE for proteoforms, ^[23]^ high sensitivity of CZE-MS for proteoform detection, ^[23–25]^ and high orthogonality of LC and CZE for biomolecule separations. ^[21,26]^ The features of CZE-MS/MS for TDP have been systematically reviewed recently. ^[27,28]^ The list of identified proteoforms is shown in **Supporting Information II**.

**Figure 2.**
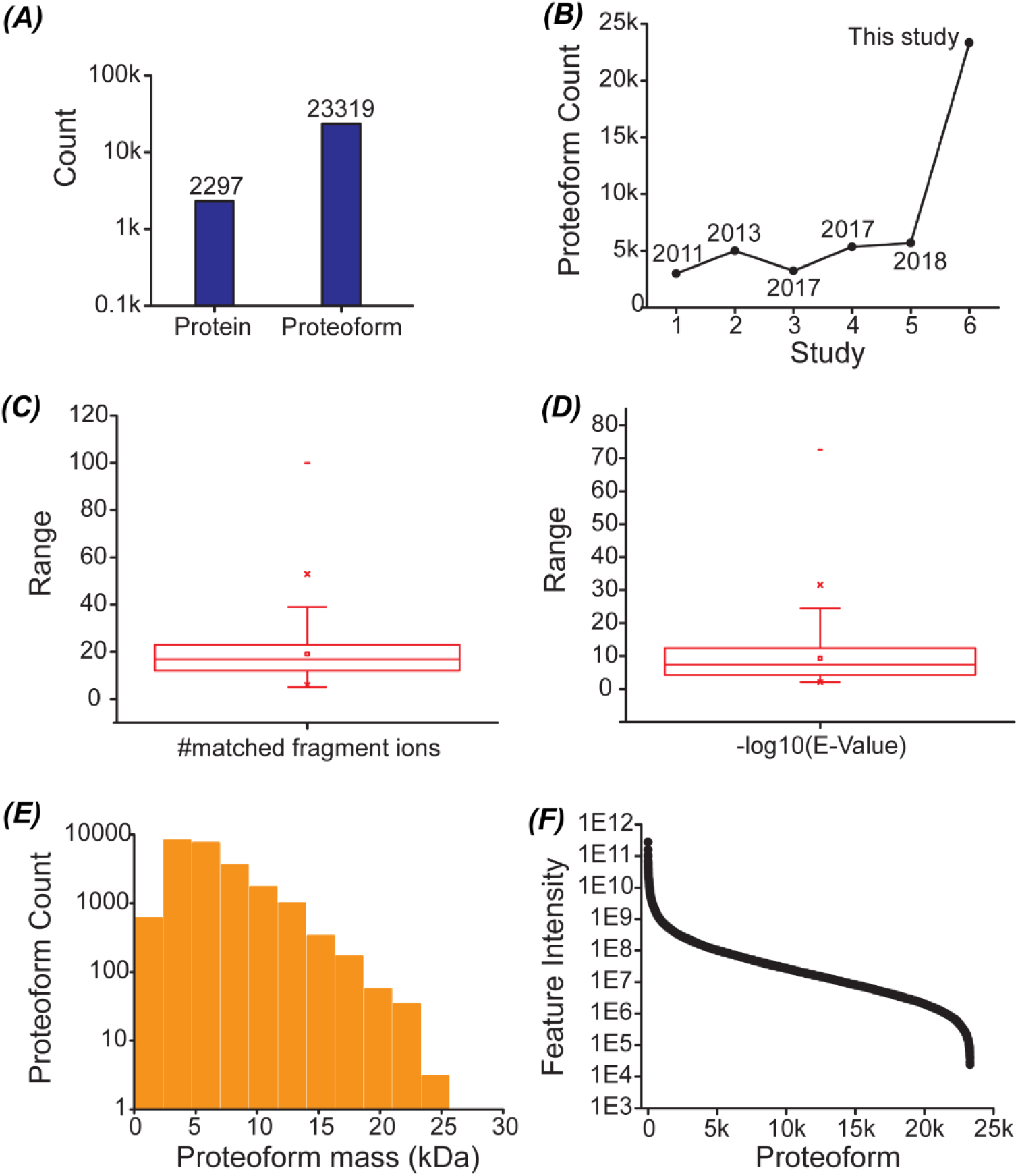
Summary of proteoform identifications from the study. (A) Protein and proteoform identifications with a 1% proteoform FDR. (B) Learning curve of the number of proteoform IDs from complex proteomes using RPLC- or CZE-based TDP. Box plots of the distribution of the number of matched fragment ions (C) and −log10 (E-value) of the identified proteoforms (D). (E) Distribution of the mass of identified proteoforms. (F) Plot of feature intensity of identified proteoforms showing the high abundance dynamic range of the proteoforms.

As shown in **Figures 2C** and **2D**, the proteoforms were identified with high confidence. The average number of matched fragment ions per proteoform is nearly 20 and the average E-value of the identified proteoforms is about 1E-10. We identified 2754 proteoforms with masses in a range of 10-26 kDa and the majority of identified proteoforms had masses smaller than 10 kDa, **Figure 2E**. The intensity of identified proteoforms spanned seven orders of magnitude, **Figure 2F**, indicating the wide concentration dynamic range of proteoforms in the human cell samples.

Protein PTMs modulate their biological function. For example, protein N-terminal acetylation influences the stability, folding, binding, and subcellular targeting of proteins.^[29]^ Protein phosphorylation is well known for regulating cell signaling, gene expression, and differentiation.^[30]^ Protein methylation plays important roles in modulating transcription.^[31]^ This large-scale TDP study identified 4872 proteoforms with N-terminal acetylation (+42 Da mass shift), 319 proteoforms with phosphorylation [+80 Da (single phosphorylation) or +160 Da (double phosphorylation) mass shift], 321 proteoforms with methylation (+14 Da mass shift), and 241 proteoforms with oxidation (+16 Da mass shift), **Figure 3A**. TDP is powerful for the characterization of combinations of various PTMs on each proteoform. Here we identified 54 proteoforms with two phosphorylation sites, and identified 90 proteoforms with both acetylation and phosphorylation PTMs. **Figure 3B** shows the sequences and fragmentation patterns of phosphorylated and unphosphorylated proteoforms of AKAP8L (A-kinase anchor protein 8-like). AKAP8L is associated with an unfavorable prognosis in CRC based on the Human Protein Atlas (https://www.proteinatlas.org/ENSG00000011243-AKAP8L). Both the two proteoforms of AKAP8L were identified with reasonably high confidence and the phosphorylation site was localized to the S601. The S601 phosphorylation on AKAP8L was only reported in a liver phosphoproteome study using the bottom-up strategy ^[32]^ and has not been reported in the two CRC cell lines studied here according to the PhosphoSitePlus (version 6.5.9.3). ^[33]^ More studies are needed to reveal the function of phosphorylated AKAP8L at S601 in CRC.

**Figure 3.**
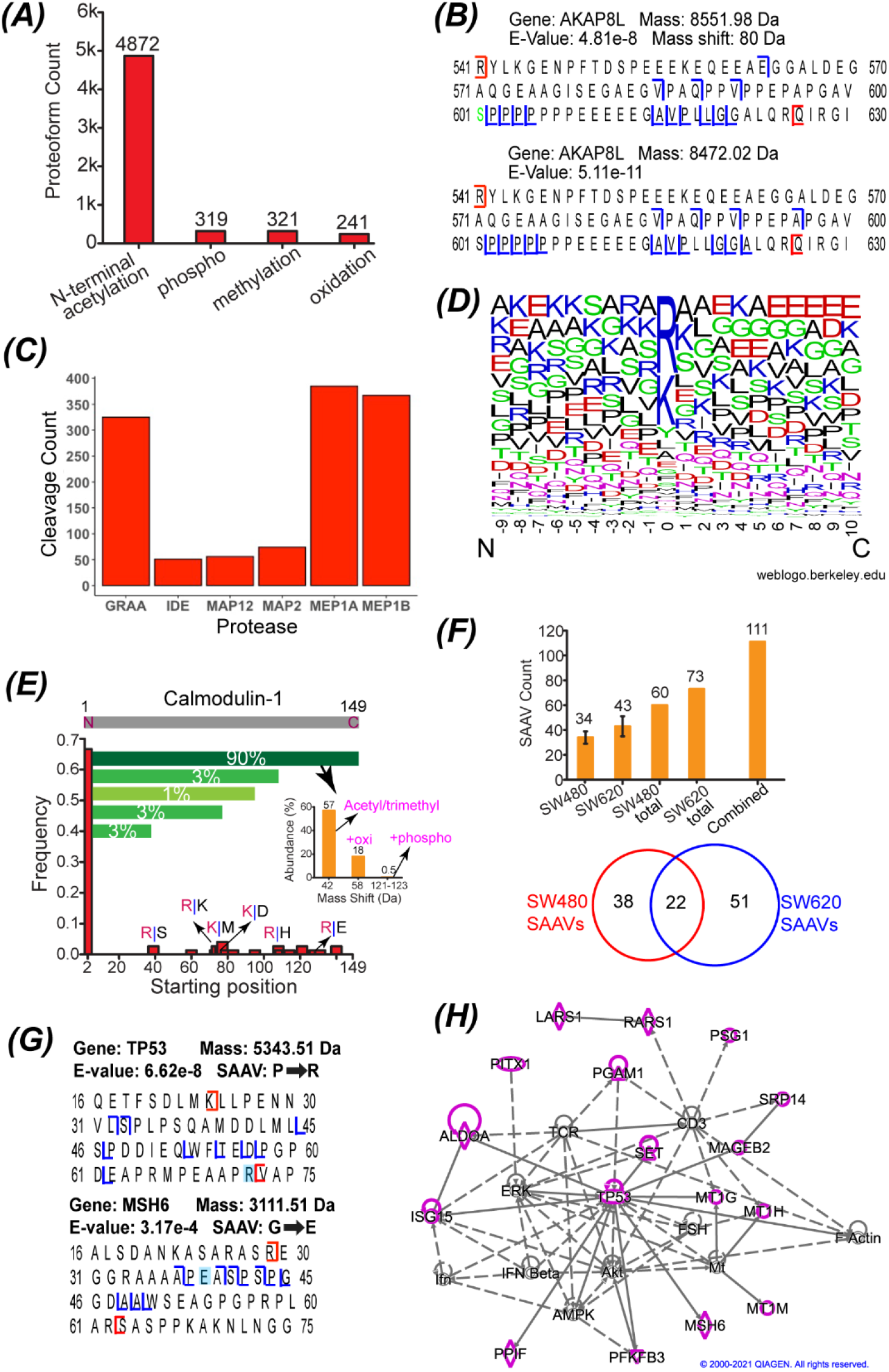
Deep analyses of the identified proteoforms from CRC cells. (A) Proteoforms with various PTMs, including N-terminal acetylation, phosphorylation, methylation, and oxidation. (B) Sequences and fragmentation patterns of two proteoforms (phosphorylated and un-phosphorylated) of AKAP8L. (C) Summary of the TopFINDer analysis result regarding the enzymatic cleavages by proteases (P-value<0.05). GRAA: Granzyme A; IDE: Insulin-degrading enzyme; MAP12: Methionine aminopeptidase 1D, mitochondrial; MAP2: Methionine aminopeptidase 2; MEP1A: Meprin A subunit alpha; MEP1B: Meprin A subunit beta. (D) Weblogo data of the amino acid residues surrounding the truncation sites (position 0) of N-terminally truncated proteoforms identified in the study. The WebLogo tool (https://weblogo.berkeley.edu/) was used to create the figure. (E) Summary of all the identified proteoforms of calmodulin-1 regarding starting positions, relative abundance based on the number of PrSMs, and PTMs. (F) The number of proteoforms containing SAAVs identified from the SW480 and SW620 cells and the overlap of those proteoforms. The error bars in the figure represent the standard deviations of proteoforms from triplicate measurements. (G) Sequences and fragmentation patterns of two proteoforms containing SAAVs. (H) SAAVs containing proteoforms correspond to many genes (highlighted in purple) that are involved in a cancer related network according to the IPA analysis. The diamond, triangle, oval, and circle shapes represent proteins belonging to enzyme, phosphatase/kinase, transcription regulator and other, respectively. The solid and dotted lines represent direct and indirect interactions.

We noted that both proteoforms of AKAP8L in **Figure 3B** were truncated at N and C termini. Actually, in this study we identified many truncated proteoforms. Some of the truncations could be due to the enzymatic processing in the cells. To test this hypothesis, we utilized the TopFINDer tool to seek clues of enzymatic cleavages by analyzing the truncated proteoforms. ^[34]^ We determined protein cleavage activities of six enzymes with p-value better than 0.05, **Figure 3C**, including GRAA (Granzyme A, cleaves after K or R), ^[35,36]^ IDE (Insulin-degrading enzyme, preferentially cleaves hydrophobic and basic residues),^[37]^ MAP12 (Methionine aminopeptidase 1D, mitochondrial, removes the N-terminal M from proteins), MAP2 (Methionine aminopeptidase 2, removes the N-terminal M from proteins), MEP1A (Meprin A subunit alpha, hydrolyze proteins preferentially on hydrophobic residues), and MEP1B (Meprin A subunit beta, has a preference for acidic amino acids after the cleavage site) ^[38]^. Additionally, manual examinations of the identified proteoforms confirmed the presence of some complementary proteoform sequences to the cleaved sequences, which further improved the confidence of the enzymatic activity results. The MAP12 protein is overexpressed in CRC cells and tumors compared to normal samples and may play some important roles in CRC tumorigenesis.^[39]^ We also analyzed over 6,000 N-terminally truncated proteoforms regarding the amino acids surrounding the truncation sites, **Figure 3D**. Basic amino acid residues (K and R) are highly enriched at the truncation sites (position 0), which may due to the GRAA and IDE activities. Trypsinogen has been found in various CRC cell lines and tumors, and the proteoform truncations at the K and R positions may also attribute to the activity of human trypsin in CRC cells.^[40,41]^ It has been demonstrated that trypsinogen has higher abundance in metastatic CRC cell lines (e.g. SW620) than non-metastatic CRC cell lines (e.g., SW480), suggesting the potential roles of trypsinogen in CRC invasion and metastasis.^[40]^ We further analyzed the N-terminally truncated proteoforms identified from SW480 and SW620 cell lines in our SEC-CZE-MS/MS data produced in this study. We discovered that 71±0.6% (n=3) of those truncated proteoforms in SW620 cells were cleaved at K or R and the percentage was statistically significantly (t-test p-value 0.03) higher than that in SW480 cells (67±2%, n=3).

One important value of TDP is its capability for delineation of various proteoforms from the same gene (proteoform family).^[42]^ **Figure 3E** shows one example of Calmodulin-1 (*CALM1*) proteoform family. Calmodulin-1 modulates many enzymes (kinases and phosphatases), ion channels, and many other proteins by calcium-binding. We identified 75 proteoforms of *CALM1*. Nearly 70% of those proteoforms start at the position 2 with the N-terminal methionine removal. Various truncated proteoforms, for example, with the starting positions around 40, 60, 80 and 120, were identified in a much lower frequency. Many of those truncated proteoforms were cleaved at the basic amino acid residues (K or R). The number of proteoform spectrum matches (PrSMs) can be used to roughly estimate the relative abundance of proteoforms.^[21,43]^ For the *CALM1* proteoforms starting from position 2, about 90% of the corresponding PrSMs match to proteoforms covering the whole protein sequence (2-149), called intact proteoforms. The PrSMs corresponding to other C-terminally truncated proteoforms only account for 3% or lower. The intact proteoforms have various PTMs, including acetylation/trimethylation, oxidation, and phosphorylation. The intact proteoforms of *CALM1* with a 42-Da mass shift (acetylation/trimethylation) are the most abundant forms; intact proteoforms with additional oxidation (a 58-Da mass shift) or phosphorylation (a 122-Da mass shift) have much lower abundance according to the number of PrSMs of those proteoforms.

Cancers result from gene mutations, which produce proteoforms containing amino acid variants (AAVs). Although transcriptomic analysis can provide ample information about gene mutations and possible AAVs on proteins, it is valuable to detect proteoforms containing AAVs directly because gene expression can be regulated post-transcriptionally. Bottom-up proteomics has been used for the identification of peptides containing single AAVs (SAAVs) from cancer cells.^[44,45]^ The Kelleher group reported the identification of 10 proteoforms containing SAAVs from breast tumor xenografts in one TDP study.^[46]^ Here we identified 111 proteoforms containing SAAVs of 82 genes from the SW480 and SW620 cell lines with a proteogenomic approach with a 5% proteoform-level FDR, representing one order of magnitude improvement in the number of identified proteoforms containing SAAVs compared to previous studies, **Figure 3F**. The transcriptomic variants based on the available RNA-Seq data were incorporated into the protein database for the identification of proteoforms containing SAAVs using TopPG, a recently developed bioinformatics tool.^[47]^ We also manually inspected the MS/MS spectra of proteoforms containing the SAAV sites to ensure high-confidence identifications. We identified more proteoforms containing SAAVs from metastatic SW620 cells than the non-metastatic SW480 cells (73 *vs*. 60). Only 20% of the 111 proteoforms were identified from both cell lines, suggesting drastic differences between the two cell lines regarding SAAVs profile.

**Figure 3G** shows the sequences and fragmentation patterns of two examples of proteoforms containing SAAVs. TP53 is an important tumor suppressor and it has been closely related to CRC development. We identified one proteoform containing the AAV at position 72 (P^→^ R) due to the codon 72 polymorphism. Studies have shown the functional differences of the P72 and R72 proteoforms of TP53. ^[48–50]^ For example, the R72 proteoform does a markedly better job of inducing apoptosis compared to the P72 proteoform. ^[48,49]^ In another study, the results indicated that the expression of P72 proteoform increased CRC metastasis, and R72 proteoform does not exist in the non-metastatic CRC cell line (SW480) based on the nucleic-acid data. ^[50]^ Interestingly, we only identified the R72 proteoform of TP53 in the SW620 cell line, not in the SW480 cell line, from the top-down MS data. *MSH6* is one of the DNA mismatch repair genes and its mutations play a crucial role in Lynch syndrome, which is an inherited form of CRC. We identified one MSH6 proteoform containing the SAAV due to polymorphism at position 39 (G^→^ E). The G39 has been associated with an increased risk of CRC according to the nucleic-acid data. ^[51]^ We identified G39 proteoforms of MSH6 in both SW480 and SW620 cells, but identified the E39 proteoform only in the SW480 cells, not in the SW620 cells.

For the proteoforms containing SAAVs, we further performed QIAGEN Ingenuity Pathway Analysis (IPA) analyses of the corresponding 82 genes. We revealed that 75 of those genes are associated with tumorigenesis of tissue (p-value: 0.0001), and three genes (MSH6, PITX1 and TP53) relate to the development of colon tumor (p-value: 0.002). Five of the genes related to tumorigenesis of tissue (AURKA, EIF5A, PFKFB3, POLE4, and TP53) are targets of cancer drugs. We further performed IPA network analysis and revealed that 17 out of the 82 genes are involved in a cancer-related network (network score 36), **Figure 3H**, suggesting their crucial roles in cancer and development. The 17 genes are highlighted in purple and those proteins belong to several different families, including enzyme (diamond shape, *LARS1*, *PARS1*, *ALDOA*, *MSH6*, and *PPIF*), phosphatase/kinase (triangle shape, *PGAM1*, *SET*, and *PFKFB3*), transcription regulator (oval shape, *TP53* and *PITX1*), and others (circle shape, *PSG1*, *SRP14*, *MAGEB2*, *MT1G*, *MT1H*, *MT1M*, and *ISG15*). Nine of those highlighted proteins have direct (solid line) or indirect (dotted line) interactions with TP53.

We further carried out the first quantitative TDP study of a pair of metastatic (SW620) and non-metastatic (SW480) human CRC cell lines. The cell lysates of SW480 and SW620 cells were fractionated by SEC and each fraction was analyzed by CZE-MS/MS in technical triplicate. After database search with the TopPIC and label-free quantification analysis using TopDiff (version 1.3.4), available in the TopPIC suite, we quantified nearly 1500 proteoforms and 460 proteoforms of 248 proteins showed statistically significant differences in abundance between the two cell lines (FDR<0.05), **Figure 4A**. 244 proteoforms of 152 proteins had higher abundance in the SW480 cell line and 216 proteoforms of 132 proteins had higher expression in the SW620 cell lines. Comparing the proteoforms with higher expression in SW480 cells with that in SW620 cells revealed that 36 genes were overlapped, suggesting that for those 36 genes, different proteoforms of the same gene had completely different expression patterns in the two cell lines. **Figure 4B** shows two differentially expressed proteoforms of one of those 36 genes, *DAP* (Death-associated protein 1). It has been reported that DAP modulates cell death and correlates with the clinical outcome of CRC patients.^[52]^ Interestingly, we revealed that one phosphorylated proteoform of DAP (∼7,607 Da, phosphorylation site S51 or T56) had higher abundance in SW480 cells and another phosphorylated proteoform (∼4,605 Da, phosphorylation site S51) showed higher expression in SW620 cells. Both the S51 and T56 are known to be phosphorylated according to PhosphoSitePlus, with S51 being the most common phosphorylation site. Those two proteoforms could have the same or closely localized phosphorylation sites, and they were cleaved at the basic amino acid residue, arginine (R), at both N and C termini. The data highlights the value of TDP for quantitative characterization of proteins in a proteoform-specific manner. We noted that the differentially expressed proteoforms in this study include phosphorylated proteoforms of several important genes related to CRC, i.e., *RALY*,^[53]^ *NPM1*,^[54]^ *DAP*,^[52]^ and *HDGF*,^[55]^ **Table S1**. The functions of phosphorylated forms of those four proteins in modulating CRC development are still unclear. However, the differential expressions of those phosphorylated proteoforms in the metastatic and non-metastatic CRC cells suggest their potential roles in regulating CRC metastasis.

**Figure 4.**
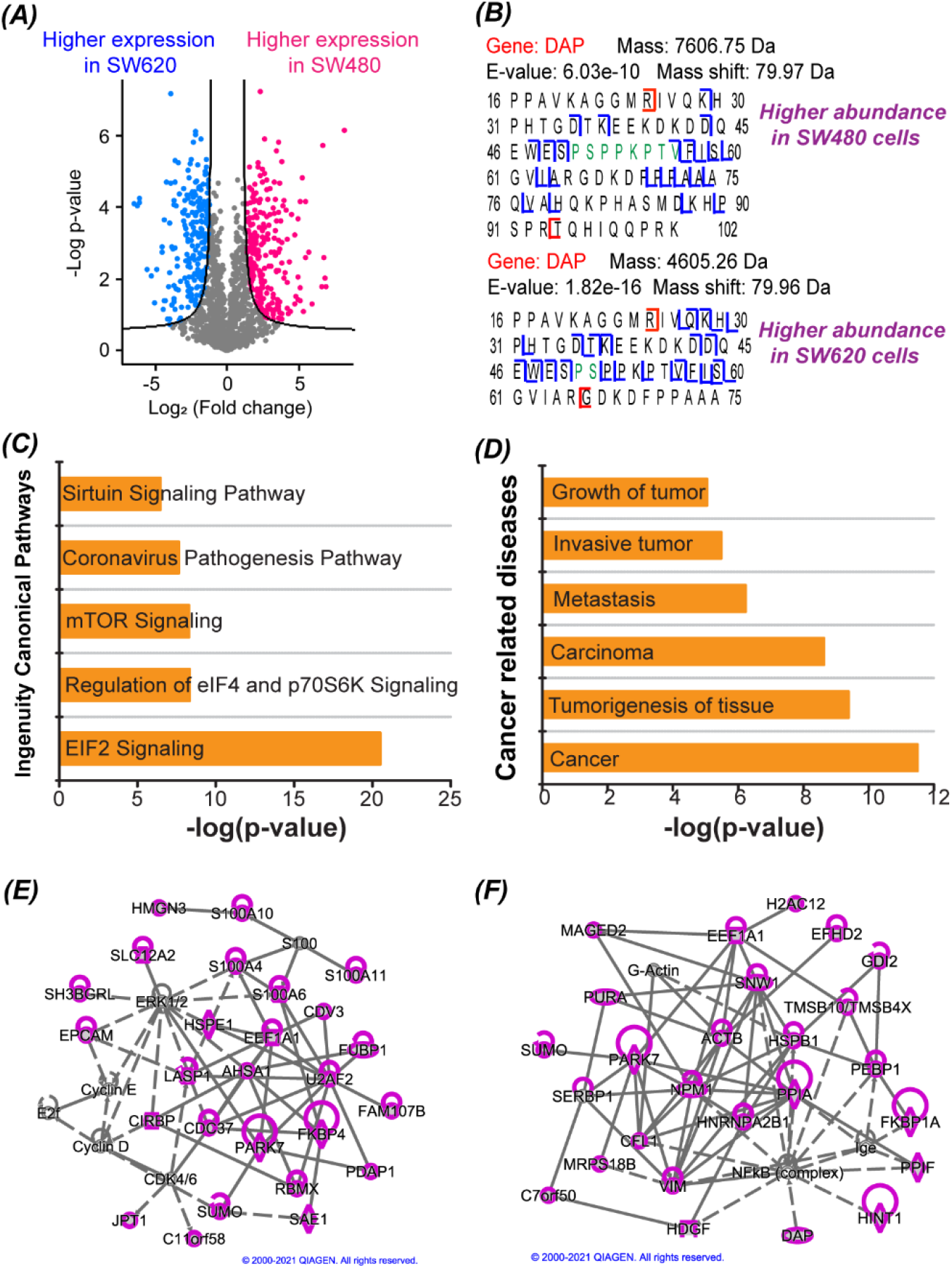
Summary of the quantitative TDP data of SW480 and SW620 cells. (A) Volcano plot showing the differentially expressed proteoforms between SW480 and SW620 cells. Pink dots and blue dots represent proteoforms having statistically significantly higher abundance in SW480 and in SW620, respectively. The Perseus software was used for performing the t-test and generating the Volcano plot with the following settings (S0=1 and FDR = 0.05). ^[56]^ (B) Sequences and fragmentation patterns of two phosphorylated proteoforms of the gene *DAP*. One has higher abundance in SW480 cells and the other has higher expression in SW620 cells. (C) The top 5 Ingenuity canonical pathways for the differentially expressed genes at the proteoform level according to the IPA analysis. (D) Some of the cancer related diseases that the differentially expressed genes are involved in according to the IPA analysis. Proteoforms with higher abundance in SW480 cells (E) or higher abundance in SW620 cells (F) correspond to genes that are involved in cancer-related networks with high scores. Those genes are highlighted in purple. The diamond, oval, hexagon, trapezium, square, and circle shapes represent enzyme, transcription regulator, translation regulator, transporter, growth factor, and other. The solid and dotted lines represent direct and indirect interactions.

We then performed IPA analyses of the genes of those differentially expressed proteoforms between SW480 and SW620 cells. The top five pathways that those genes are involved in are EIF2 signaling, regulation of eIF4 and p70S6K signaling, mTOR signaling, coronavirus pathogenesis pathway, and sirtuin signaling pathway, **Figure 4C**. Those genes are heavily involved in cancer-related diseases, for example, tumorigenesis of tissue and metastasis, **Figure 4D**. Five of those proteins (EIF4E, EPCAM, FKBP1A, GAA, and HSP90AB1) are drug targets. IPA network analyses revealed that 26 proteins (highlighted in purple) whose proteoforms showed higher abundance in SW480 compared to SW620 were involved in a cancer-related network (score 51), **Figure 4E**. Those proteins belong to several families, including enzyme (diamond shape, e.g., PARK7 and FKBP4), transcription regulator (oval shape, e.g., FUBP1), translation regulator (hexagon shape, e.g., CIRBP and EEF1A1), transporter (trapezium shape, e.g., SLC12A2 and LASP1), and other (circle shape, e.g., EPCAM and JPT1). Most of those proteins have direct (solid line) and indirect (dotted line) interactions with one another. We also carried out network analysis for the proteins whose proteoforms had higher expression in SW620 cells, and observed high-scores for cancer-related networks. **Figure 4F** shows one cancer-related network (score 54), and 26 of those proteins are involved in the network (highlighted in purple). Those proteins include several CRC-related important proteins, NPM1 (oval shape, transcription regulator, located in nucleus), DAP (transcription regulator, located in cytoplasm), and HDGF (square shape, growth factor, located in extracellular space). NPM1 is a crucial protein in the network and many of the highlighted proteins have direct interactions (solid line) with NPM1, for example, PARK7, VIM, and PPIA. NPM1 also has indirect interaction (dotted line) with the NFkB complex, which plays crucial roles in modulating DNA transcription and cell survival. Human NPM1 boosts the activation of NFkB according to Ingenuity relationships from the IPA analysis. Besides NPM1, several other highlighted proteins (e.g., HDGF and DAP) also have indirect interactions with the NFkB complex. For example, NFkB regulates the transcription of *HDGF*, and DAP deactivates the NFkB according to the IPA network analysis results.

In summary, we present the largest TDP dataset so far containing over 23000 proteoforms and performed the first quantitative TDP study of the pair of isogenic CRC cell lines (SW480 and SW620), revealing drastic differences in proteoform profiles between the metastatic and non-metastatic CRC cell lines. Many differentially expressed genes at the proteoform level with diversified functions are closely related to cancer according to the IPA analysis. We observed remarkable enzymatic processing of proteins in the two CRC cell lines, especially the cleavages at basic amino acid residues (K and R), which is potentially due to the activities of GRAA, IDE, and trypsinogen. We produced the largest dataset of proteoforms containing SAAVs and detected significant difference in the profile of proteoforms containing SAAVs between the metastatic and non-metastatic CRC cell lines. Overall, this study has taken TDP to the next level regarding proteome coverage and is a milestone in pursuing the delineation of human proteome in a proteoform-specific manner.

## Acknowledgements

The work was funded by National Cancer Institute (NCI) through the grant R01CA247863 (Sun, Hummon, and Liu). We also thank the support from National Institute of General Medical Sciences (NIGMS) through grants R01GM125991 (Sun and Liu) and R01GM118470 (Liu and Sun). Sun also thanks the support from the National Science Foundation (CAREER Award, Grant DBI1846913). We thank MSU AgBioResearch and Michigan State University for the access to QIAGEN Ingenuity Pathway Analysis (IPA) platform.

## Author contribution statements

E.N.M. performed the experiments for proteoform identifications. T.X. performed the experiment for proteoform quantification. W.C. carried out all the database search using TopPIC for proteoform ID and quantification. E.N.M., T.X., and W.C. worked together for data analysis and made the first draft of the manuscript. N.C.B. did all the cell culture and initial sample preparation of SW480 and SW620 cells. S.M.N. performed the LC fractionations. A.B.H., X.L., and L.S. conceived the original idea. X.L. supervised the database search part of the project. L.S. supervised the project. All authors provided comments and contributed to the final manuscript.

## Competing interests

The authors declare no competing interests.

## Notes

### Competing Interest Statement

The authors have declared no competing interest.

## References

1. Schmitt, M. & Greten, F. R. The inflammatory pathogenesis of colorectal cancer. Nat. Rev. Immunol. 21, 653–667 (2021).

2. Rehman, S. K. et al. Colorectal Cancer Cells Enter a Diapause-like DTP State to Survive Chemotherapy. Cell 184, 226–242 (2021).

3. Markowitz, S. D. & Bertagnolli, M. M. Molecular Basis of Colorectal Cancer. N. Engl. J. Med. 361, 2449–2460 (2009).

4. Schunter, A. J., Yue, X. & Hummon, A. B. Phosphoproteomics of colon cancer metastasis: comparative mass spectrometric analysis of the isogenic primary and metastatic cell lines SW480 and SW620. Anal. Bioanal. Chem. 409, 1749–1763 (2017).

5. Zhang, B. Clinical potential of mass spectrometry-based proteogenomics. Nat. Rev. Clin. Oncol. 16, 256–268 (2019).

6. Xu, L., Wang, R., Ziegelbauer, J., Wu, W. W., Shen, R. F., Juhl, H., Zhang, Y., Pelosof, L. & Rosenberg, A. S. Transcriptome analysis of human colorectal cancer biopsies reveals extensive expression correlations among genes related to cell proliferation, lipid metabolism, immune response and collagen catabolism. Oncotarget. 8, 74703–74719 (2017).

7. Huo, T., Canepa, R., Sura, A., Modave, F. & Gong, Y. Colorectal cancer stages transcriptome analysis. PLoS One 12, e0188697 (2017).

8. Zhang, B. et al. Proteogenomic characterization of human colon and rectal cancer. Nature 513, 382–387 (2014).

9. Besson, D. et al. A quantitative proteomic approach of the different stages of colorectal cancer establishes OLFM4 as a new nonmetastatic tumor marker. Mol. Cell. Proteomics 10, M111.009712 (2011).

10. Ghosh, D. et al. Identification of Key Players for Colorectal Cancer Metastasis by iTRAQ Quantitative Proteomics Profiling of Isogenic SW480 and SW620 Cell Lines. J. Proteome Res. 10, 4373–4387 (2011).

11. Smith, L. M. et al. Proteoform: a single term describing protein complexity. Nat Methods 10, 186–187 (2013).

12. Smith L. M. & Kelleher, N. L. Proteoforms as the next proteomics currency. Science 359, 1106–1107 (2018).

13. Toby, T. K., Fornelli, L. & Kelleher, N. L. Progress in Top-Down Proteomics and the Analysis of Proteoforms. Annu. Rev. Anal. Chem. 9, 499–519 (2016).

14. Ntai, I. et al. Precise characterization of KRAS4b proteoforms in human colorectal cells and tumors reveals mutation/modification cross-talk. Proc Natl Acad Sci U S A. 115, 4140–4145.

15. Kou, Q., Xun, L. & Liu, X. TopPIC: a software tool for top-down mass spectrometry-based proteoform identification and characterization. Bioinformatics 32, 3495–3497 (2016).

16. Smith, L. et al. The Human Proteoform Project: A Plan to Define the Human Proteome. Preprints 2020100368 doi: 10.20944/preprints202010.0368.v1 (2020).

17. Aebersold, R. et al. How many human proteoforms are there? Nat. Chem. Biol. 14, 206–214 (2018).

18. Tran, J. C. et al. Mapping intact protein isoforms in discovery mode using top-down proteomics. Nature 480, 254–258 (2011).

19. Catherman, A. C. et al. Large-scale top-down proteomics of the human proteome: membrane proteins, mitochondria, and senescence. Mol. Cell. Proteomics 12, 3465–3473 (2013).

20. Anderson, L. C. et al. Identification and Characterization of Human Proteoforms by Top-Down LC-21 Tesla FT-ICR Mass Spectrometry. J. Proteome Res. 16, 1087–1096 (2017).

21. McCool, E. N. et al. Deep Top-Down Proteomics Using Capillary Zone Electrophoresis-Tandem Mass Spectrometry: Identification of 5700 Proteoforms from the Escherichia coli Proteome. Anal Chem. 90, 5529–5533 (2018).

22. Cai, W. et al. Top-Down Proteomics of Large Proteins up to 223 kDa Enabled by Serial Size Exclusion Chromatography Strategy. Anal. Chem. 89, 5467–5475 (2017).

23. Lubeckyj, R. A., Basharat, A. R., Shen, X., Liu, X. & Sun, L. Large-Scale Qualitative and Quantitative Top-Down Proteomics Using Capillary Zone Electrophoresis-Electrospray Ionization-Tandem Mass Spectrometry with Nanograms of Proteome Samples. J. Am. Soc. Mass Spectrom. 30, 1435–1445 (2019).

24. McCool, E. N. & Sun, L. Comparing nanoflow reversed-phase liquid chromatography-tandem mass spectrometry and capillary zone electrophoresis-tandem mass spectrometry for top-down proteomics. Se Pu 37, 878–886 (2019).

25. Han, X. et al. In-line separation by capillary electrophoresis prior to analysis by top-down mass spectrometry enables sensitive characterization of protein complexes. J. Proteome Res. 13, 6078–6086 (2014).

26. Yang, Z., Shen, X., Chen, D. & Sun, L. Improved Nanoflow RPLC-CZE-MS/MS System with High Peak Capacity and Sensitivity for Nanogram Bottom-up Proteomics. J Proteome Res. 18, 4046–4054 (2019).

27. Chen, D. et al. Recent advances (2019-2021) of capillary electrophoresis-mass spectrometry for multilevel proteomics. Mass Spectrom. Rev. doi: 10.1002/mas.21714 (2021).

28. Gomes, F. P. & Yates, J. R. 3^rd^. Recent trends of capillary electrophoresis-mass spectrometry in proteomics research. Mass Spectrom. Rev. 38, 445–460 (2019).

29. Ree, R., Varland, S. & Arnesen, T. Spotlight on protein N-terminal acetylation. Exp. Mol. Med. 50, 1–13 (2018).

30. Kalume, D. E., Molina, H. & Pandey, A. Tackling the phosphoproteome: tools and strategies. Curr. Opin. Chem. Biol. 7, 64–69 (2003).

31. Lee, D. Y., Teyssier, C., Strahl, B. D. & Stallcup, M. R. Role of protein methylation in regulation of transcription. Endocr. Rev. 26, 147–170 (2005).

32. Bian, Y. et al. An enzyme assisted RP-RPLC approach for in-depth analysis of human liver phosphoproteome. J. Proteomics 96, 253–262 (2014).

33. Hornbeck, P. V., Zhang, B., Murray, B., Kornhauser, J. M., Latham, V. & Skrzypek, E. PhosphoSitePlus, 2014: mutations, PTMs and recalibrations. Nucleic Acids Res. 43, D512–D520 (2015).

34. Fortelny, N., Yang, S., Pavlidis, P., Lange, P. F. & Overall, C. M. Proteome TopFIND 3.0 with TopFINDer and PathFINDer: database and analysis tools for the association of protein termini to pre- and post-translational events. Nucleic Acids Res. 43, D290–D297 (2015).

35. Zhou, Z. et al. Granzyme A from cytotoxic lymphocytes cleaves GSDMB to trigger pyroptosis in target cells. Science 368, eaaz7548 (2020).

36. Hink-Schauer, C., Estébanez-Perpiñá, E., Kurschus, F. C., Bode, W. & Jenne, D. E. Crystal structure of the apoptosis-inducing human granzyme A dimer. Nat. Struct. Biol. 10, 535–540 (2003).

37. Song, E. S. et al. Analysis of the subsite specificity of rat insulysin using fluorogenic peptide substrates. J. Biol. Chem. 276, 1152–1155 (2001).

38. Becker-Pauly, C. et al. Proteomic analyses reveal an acidic prime side specificity for the astacin metalloprotease family reflected by physiological substrates. Mol. Cell. Proteomics 10, M111.009233 (2011).

39. Leszczyniecka, M. et al. MAP1D, a novel methionine aminopeptidase family member is overexpressed in colon cancer. Oncogene 25, 3471–3478 (2006).

40. Williams, S. J., Gotley, D. C. & Antalis, T. M. Human trypsinogen in colorectal cancer. Int. J. Cancer 93, 67–73 (2001).

41. Oyama. K. et al. Trypsinogen expression in colorectal cancers. Int. J. Mol. Med. 6, 543–548 (2000).

42. Dai, Y. et al. Constructing Human Proteoform Families Using Intact-Mass and Top-Down Proteomics with a Multi-Protease Global Post-Translational Modification Discovery Database. J. Proteome Res. 18, 3671–3680 (2019).

43. Geis-Asteggiante, L., Ostrand-Rosenberg, S., Fenselau, C. & Edwards, N. J. Evaluation of Spectral Counting for Relative Quantitation of Proteoforms in Top-Down Proteomics. Anal. Chem. 88, 10900–10907 (2016).

44. Tan, Z. et al. Comprehensive Detection of Single Amino Acid Variants and Evaluation of Their Deleterious Potential in a PANC-1 Cell Line. J. Proteome Res. 19, 1635–1646 (2020).

45. Song, C. et al. Large-scale quantification of single amino-acid variations by a variation-associated database search strategy. J. Proteome Res. 13, 241–248 (2014).

46. Ntai, I. et al. Integrated Bottom-Up and Top-Down Proteomics of Patient-Derived Breast Tumor Xenografts. Mol. Cell. Proteomics 15, 45–46 (2016).

47. Chen, W. & Liu, X. Proteoform Identification by Combining RNA-Seq and Top-Down Mass Spectrometry. J. Proteome Res. 20, 261–269 (2021).

48. Dumont, P., Leu, J. I., Pietra 3rd, A. C. D., George, D. L. & Murphy, M. The codon 72 polymorphic variants of p53 have markedly different apoptotic potential. Nat. Genet. 33, 357–365 (2003).

49. Jeong, B., Hu, W., Belyi, V., Rabadan, R. & Levine, A. J. Differential levels of transcription of p53-regulated genes by the arginine/proline polymorphism: p53 with arginine at codon 72 favors apoptosis. FASEB J. 24, 1347–1353 (2010).

50. Katkoori, V. R. et al. Functional consequence of the p53 codon 72 polymorphism in colorectal cancer. Oncotarget. 8, 76574–76586 (2017).

51. Zelga, P., Przybyłowska-Sygut, K., Zelga, M., Dziki, A. & Majsterek, I. Polymorphism of Gly39Glu (c.116G>A) hMSH6 is associated with sporadic colorectal cancer development in the Polish population: Preliminary results. Adv. Clin. Exp. Med. 26, 1425–1429 (2017).

52. Jia, Y. et al. Death associated protein 1 is correlated with the clinical outcome of patients with colorectal cancer and has a role in the regulation of cell death. Oncol. Rep. 31, 175–182 (2014).

53. Sun, L. et al. RNA-binding protein RALY reprogrammes mitochondrial metabolism via mediating miRNA processing in colorectal cancer. Gut 70, 1698–1712 (2021).

54. Grisendi, S., Mecucci, C., Falini, B. & Pandolfi, P. P. Nucleophosmin and cancer. Nat. Rev. Cancer. 6, 493–505 (2006).

55. Sun, B., Gu, X., Chen, Z. & Xiang, J. MiR-610 inhibits cell proliferation and invasion in colorectal cancer by repressing hepatoma-derived growth factor. Am. J. Cancer Res. 5, 3635–3644 (2015).

56. Tyanova S. et al. The Perseus computational platform for comprehensive analysis of (prote)omics data. Nat. Methods 13, 731–740 (2016).

