## supporting information I for "Qualitative and quantitative top-down proteomics of human colorectal cancer cell lines identified 23000 proteoforms and revealed drastic proteoform-level differences between metastatic and non-metastatic cancer cells"

Amanda B. Hummon

Xiaowen Liu

Liangliang Sun

**Table of Contents**


| | Page |
|---|---|
| **Experimental Procedures** | 3-9 |
| **Table S1** | 10 |
| **References** | 11-12 |

### Experimental Procedures

#### *Materials and Reagents*

MS-grade water, acetonitrile (ACN), methanol (MeOH), formic acid (FA) and HPLC-grade acetic acid (AA) were purchased from Fisher Scientific (Pittsburgh, PA). Ammonium bicarbonate (NH4HCO3), urea, dithiothreitol (DTT), iodoacetamide (IAA) and 3-(trimethoxysilyl)propyl methacrylate were from Sigma-Aldrich (St. Louis, MO). Hydrofluoric acid (HF, 48-51% solution in water) and acrylamide were purchased from Acros Organics (NJ, USA). Fused silica capillaries (50 µm i.d./360 µm o.d.) were purchased from Polymicro Technologies (Phoenix, AZ). Complete, mini protease inhibitor cocktail (EASYpacks) was from Roche (Indianapolis, IN).

#### *Sample Preparation*

SW480 and SW620 cells were both purchased from ATCC (Manassas, VA) and were cultured in RPMI 1640 cell culture medium (Life Technologies Corporation, Grand Island, NY) supplemented with 10% fetal bovine serum (Thermo Scientific, Gaithersburg, MD) and 2mM L-glutamine (Invitrogen, San Diego, CA). The cells were incubated at 37°C with 5% CO2 and were passaged every 3-4 days. Both cell lines were last verified by Short Tandem Repeat (STR) sequencing in 2016 and were used within two months after resuscitation from frozen aliquots at -80°C.

Upon growing to confluency, cells were harvested and cleansed of remaining cell culture medium via subsequent washing with HPLC grade water (Fisher Scientific, Pittsburgh, PA) and centrifugation for 5-minute intervals at 15000 × g until supernatant was clear. Proteins were then extracted using mammalian cell lysis buffer according to the literature.[1] Cell lysis buffer consisted of 8 M urea, 50 mM Tris (pH 8.2), 1 mM β-glycerophosphate, 1 mM phenylmethylsulfonyl fluoride, 75 mM sodium chloride, 1 mM sodium fluoride, 1 mM sodium orthovanadate, 10 mM sodium pyrophosphate, and one protease inhibitor cocktail. The reagents for cell lysis buffer were purchased from Sigma-Aldrich and complete EDTA-free protease inhibitor cocktail tablet was purchased from Roche. Lysis buffer was added to the harvested cells which then underwent sonication on ice three times for 1-minute intervals at 15% amplitude. The resulting extracted proteins were then clarified of cellular debris by centrifugation at 15,000 rpm for 10 minutes. Proteins were quantified using a bicinchoninic acid (BCA) protein assay (Thermo Scientific Pierce, Rockford, IL) and then stored at -80°C until preparation for MS analysis.

SW480 and SW620 proteins were denatured at 37 °C for 30 minutes, reduced at 37 °C for 30 minutes using DTT, and then alkylated at room temperature in the dark for 20 minutes using IAA. The excess IAA were quenched by adding DTT and reacting for 5 min at room temperature.

For the first experiment, 200 μg of proteins from SW480 and SW620 cells were reduced, alkylated, and acidified, followed by RPLC fractionation and CZE-MS/MS. For the second experiment, 2 mg of proteins from SW480 and SW620 cells were reduced and alkylated before fractionated by SEC-RPLC and analyzed by CZE-MS/MS. For the third experiment, 420 μg of proteins from SW480 and SW620 cells were reduced and alkylated prior to fractionation by RPLC and analyses by CZE-MS/MS. For the fourth experiment, the samples were desalted after reduction and alkylation using a C4 trap column (4×10 mm, 3 μm particles, 300 Å pore size). Specifically, 500 μg of proteins from SW480 and SW620 cells was loaded onto the column and flushed with mobile phase A (2% (v/v) ACN, 0.1% FA) for 10 minutes at a flow rate of 1 mL/min. The proteins were eluted with mobile phase B (80% ACN, 0.1% FA) for 3 minutes at flow rate of 1 mL/min. The eluates were lyophilized with a speed vacuum and redissolved in 150 μL 0.1% formic acid (FA). Then proteins from SW480 and SW620 cells were fractionated by SEC, followed by CZE-MS/MS analyses.

***Fractionation of the SW480 and SW620 proteome***

All separations were performed on a 1260 Infinity II HPLC system from Agilent (Santa Clara, CA). Detection was performed using a UV-visible detector at a wavelength of 254 nm. Data was collected and analyzed using OpenLAB software. RPLC (C4, 2.1 × 250 mm, Sepax Technologies) and SEC (4.6 × 300 mm, 500 Å pores, Agilent) were performed offline (Agilent HPLC) for prefractionation. Fractions from SW620 and SW480 from experiment 1 (13 fractions × 2 samples), experiment 2 (84 fractions × 2 samples), experiment 3 (6 fractions × 2 samples), and experiment 4 (6 fractions × 2 samples) were analyzed by CZE-MS/MS, respectively.

In experiment 1, RPLC was used for sample fractionation with a 0.25 mL/min flow rate and gradient of 0-80% mobile phase (MP) B over 90 minutes (MPA: 2% ACN, 0.1% FA in water; MPB: 80% ACN, 0.1% FA in water). Fractions were collected from 15 to 22 minutes (fraction 1) and 22 to 70 minutes (12 fractions, 4 minutes per fraction). For experiment 2, both SEC and RPLC were used for fractionation prior to CZE-MS/MS. For SEC, the flow rate was 0.35 mL/min with a 0.05% TFA mobile phase. 2 mg of proteins in 800 µL solution was fractionated by SEC. Fractions were collected from 5-8 minutes (fraction 1) and 8-12.5 minutes (3 fractions, 1.5 minutes per fraction). One RPLC run was performed for each SEC fraction with a flow rate of 0.25 mL/min and gradient of 0-80% MPB (MPA: 2% ACN, 0.1% TFA in water; MPB: 10% IPA, 0.1% TFA in ACN) over 90 minutes with a 10-minutes equilibration with 100% MPA at the beginning of the separation. Fractions were collected from 20 to 25 minutes (fraction 1) and 25 to 65 minutes (20 fractions, 2 minutes per fraction). In experiment 3, RPLC fractionation was carried out using the same mobile phases as in experiment 1, and a 90-minute gradient was used with a 10-minute equilibration with 100% MPA at the beginning of the separation. Fractions were collected from 25 to 55 minutes (fraction 1), 50 to 70 minutes (4 fractions, 5 minutes per fraction), and 70 to 95 minutes (fraction 6). In experiment 4, SEC fractionation was performed with an Agilent Bio SEC-5 column (4.6 × 300 mm, 5 µm particles, 500 Å pore size). 220 µg of SW480 and SW620 proteins (1.5 mg/mL, 75 µL×2 injections) were loaded into the SEC column and separated isocratically at the flow rate of 0.3 mL/min with 0.1% FA as mobile phase. The first fraction is collected from 5.6 to 8.6 minutes. The second to the fifth fraction was from 8.6 to 14.6 minutes with 1.5 minutes per fraction. The final fraction was collected from 14.6 to 19.0 min. In the experiments 1-4, samples were dried down and redissolved in 50 mM $NH_4HCO_3$ (pH 8.0, ~2 mg/mL) for CZE-ESI-MS/MS.

***CZE-MS/MS analysis***

CZE separation was performed using a CESI 8000 Plus CE system (Beckman Coulter). A commercialized electrokinetically pumped sheath-flow CE-MS nanospray interface (CMP Scientific Corp) was applied for online coupling the CE system and mass spectrometer.[2,3] A glass emitter (orifice size: 20~30 µm) installed on the interface was filled with sheath buffer (0.2% FA, 10% methanol) to generate electrospray at voltage of 2-2.3 kV.

A 100 cm LPA coated fused silica capillary (50 µm i.d., 360 µm o.d.) was used for CZE separation in experiments 1, 2, and 4, while a 70 cm LPA coated capillary (50 µm i.d., 360 µm o.d.) was employed for separation in experiment 3. The inner wall of the capillary was coated with LPA based on the procedure described in references [4] and [5]. One end of the capillary was etched with HF to reduce the outer diameter of the capillary to about 70-80 µm based on the procedure described in reference [6]. (Caution: use appropriate safety procedures while handling hydrofluoric acid solutions)

In experiments 1, 2, and 4, the capillary (100 cm) was loaded with 500 nL of sample. In experiment 3, the capillary (70 cm) was loaded with ~350 nL of sample. After sample loading, the capillaries were inserted into background electrolyte, containing 5% acetic acid (pH 2.4), and 30 kV voltage was applied at the sample injection end to carry out separations.

MS1 and MS2 data were collected on a Q-Exactive HF mass spectrometer (Thermo Fisher Scientific) under data-dependent acquisition (DDA) mode. The temperature of ion transfer tube was set to 320 °C and s-lens RF was 55. MS1 spectra were collected with following parameters: m/z range of 600-2000, mass resolution of 120,000 (at m/z 200), a microscan number of 3, AGC target value of 1E6, and maximum injection time of 100 ms. The top 5 most abundant precursor ions (charge state higher than 5, or charge state unassigned and intensity threshold 2E4) in the MS1 spectra were isolated with a window of 4 m/z and fragmented via HCD with NCE of 20%. The settings for MS2 spectra were resolution of 120,000 (at m/z 200), a microscan number of 3, AGC target value of 1E5, and maximum injection time of 200 ms. The dynamic exclusion was set to a duration of 30s and the isotopic peaks were excluded.

In experiments 2, 3 and 4, each LC fraction was analyzed by CZE-MS/MS in triplicate. In experiment 1, each LC fraction was analyzed by a single CZE-MS/MS run. In total, 410 MS raw files with good protein signals were produced and used for database search, including 26 MS raw files from experiment 1 (13 fractions × 2 samples), 312 MS raw files from experiment 2 (52 fractions × 2 samples × 3 replicates), 36 MS raw files from experiment 3 (6 fractions × 2 samples × 3 replicates), and 36 MS raw files from experiment 4 (6 fractions × 2 samples × 3 replicates). We need to note that we collected 84 fractions × 2 samples in the experiment 2. However, we only observed good protein signals from 52 LC fractions per sample.

***Data analysis for proteoform identification***

All RAW files generated from the CZE-MS/MS runs were analyzed with the TopPIC Suite (version 1.4.0) pipeline.[7,8] The RAW files were converted into mzML files with msconvert.[9] Then spectral deconvolution was performed with TopFD (version 1.4.0), which converts precursor and fragment isotope clusters into neutral monoisotopic masses and finds proteoform features by combining precursor isotope clusters with similar monoisotopic masses and close migration times in MS1 scans. The resulting mass spectra with monoisotopic neutral masses were stored in msalign files and the proteoform feature information was stored in text files. The human proteome database was downloaded from UniProt (UP000005640, 20350 entries, version October 23, 2019, only reviewed protein sequences were included) and concatenated with a random decoy database of the same size. Each msalign file was searched against the concatenated targe-decoy database using TopPIC (version 1.4.0). Cysteine carbamidomethylation was set as a fixed modification, and the maximum number of unexpected modifications was 1. The precursor and fragment mass error tolerances were 15 ppm. The maximum mass shift of unknown modifications was 500 Da. TopPIC reported a list of target and decoy proteoform-spectrum-matches (PrSMs) for each msalign file.

***Merging proteoform identifications***

The proteoforms identified from all msalign files were merged and filtered with a proteoform-level FDR. First, the target and decoy PrSMs reported from all the msalign files were combined and filtered with a 5% spectrum-level FDR. [10] The PrSMs were then clustered by grouping PrSMs into the same cluster if they were from the same protein and their precursor mass differences were not large than 2.2 Da. The PrSM with the best E-value was selected for each cluster and its proteoform was reported as the representative one for the cluster. The representative target and decoy proteoforms were finally filtered with a 1% proteoform-level FDR.

***Proteoform quantification***

There were 18 MS raw files from triplicate CZE-MS/MS analyses of the 6 SEC fractions for the SW480 or SW620 sample in experiment 4. The TopPIC suite pipeline reported a list of targe and decoy PrSM identifications for each raw file. Using the methods in the previous section, the PrSM identifications of the 36 MS raw files were merged and a list of proteoform identifications with a 1% proteoform-level FDR were reported. The abundance of a proteoform was computed as the sum of the proteoform abundances in the six SEC fractions, which were reported by TopFD. Proteoform identifications and their abundances were reported for each replicate using this method. Finally, TopDiff (version 1.4.0), a tool in TopPIC Suite, was used to match proteoform identifications across the three SW480 replicates and three SW620 replicates.

The quantitative results were further analyzed using Perseus software.[11] The intensities of each proteoform in triplicate CZE-MS/MS runs of SW480 and SW620 were normalized to the intensity of corresponding proteoform from the first run of SW480, converting proteoform intensity to proteoform ratio. Then, proteoform ratios of each run were divided by the corresponding median to make sure the ratios center at 1. After log2 transformation of all the data, the significantly differentially expressed proteoforms were determined by performing t-test analysis (FDR threshold: 0.05, S0: 1). The volcano plot [-log(p-value) vs. log2(fold change)] was generated in Perseus software.

***Proteogenomic analysis***

To generate sample-specific protein sequence databases with genetic variations for SW480 and SW620 cells, two RNA-Seq data sets (SRR8616059 for SW480 and SRR8615459 for SW620) [12] were downloaded from the Sequence Read Archive (SRA). The GATK pipeline [13] was employed to align short reads in the RNA-Seq data with the hg38 human genome to call single nucleotide variants (SNVs) and indels, which were further annotated using the gene-based annotation of ANNOVAR [14] (April 16, 2018). The annotated nonsynonymous SNVs and indels in exons were chosen for generating sample-specific protein sequence databases based on the basic annotation of the hg38 human genome in GENCODE [15]. Two sample-specific protein sequence databases were generated using TopPG [16] (version 1.0): one for SW480 cells and the other for SW620 cells. Each protein sequence database contained both reference protein sequences in the basic annotation of GENCODE and protein sequences with sample-specific variants. There were 74887 entries with 51485 reference sequences and 23402 sequences with variants in the database for SW480 cells and 75665 entries with 51432 reference sequences and 24233 sequences with sample-specific variants in the database for SW620 cells. The SW480 and SW620 mass spectra in experiments 3 and 4 were searched against their corresponding sample-specific database using TopPIC (version 1.4.0) with the same parameter setting in Section "Data analysis for proteoform identification". Using the methods in Section "Merging proteoform identifications," PrSMs identified in each cell line were combined and clustered, and proteoform identifications were filtered by a 5% proteoform-level FDR. Identifications with single amino acid variant (SAAV) sites were manually inspected. If a proteoform with SAAV sites contained no unexpected mass shifts or had at least three matched fragment ions between each SAAV site and the unexpected mass shift, it was reported as a confident proteoform identification with SAAV sites.

**Table S1**. Summary of the phosphorylated proteoforms with differential expression between SW480 and SW620 cells.

| Gene | Log2(proteoform intensity ratio, SW480/ SW620) | Proteoform Sequence |
|---|---|---|
| DAP | 1.53 | R.IVQKHPHTGDTKEEKDKDDQEWES(PSPPKPTV)[79.9696]FISGVIARGDKDFPPAAAQVAHQKPHASMDKHPSPR.T |
| DAP | -1.49 | R.IVQKHPHTGDTKEEKDKDDQEWES(PS)[79.9692]PPKPTVFISGVIAR.G |
| HDGF | 3.45 | R.AGDLLED(SPK)[79.9689]RPKEAENPEGEEKEAATLEVERPLPMEVEKNSTPSEPGSGRGPPQEEEEEEDEEEEATKEDAEAPGIRDHESL. |
| NPM1 | -2.67 | K.(C)[Carbamidomethylation]GSGPVHISGQHLVAVEEDAE(SE)[79.9682]DEEEEDVKLLSISGKR.S |
| RALY | -5.52 | R.TRDDGDEEGLLTH(SEEELE)[79.9695]HSQDTDADDGALQ. |
| HIST1H1B | 2.42 | M.(S)[Acetyl]ETAPAETATPA(PVEKS)[79.9702]PAKKKATK.K |
| HMGN1 | 2.01 | K.QAEVANQETKEDLPAEN(GETKTEESPAS)[159.9318]DEAGEKEAKSD. |
| HNRNPC | -3.09 | R.SAAEMYGSVTEH(PS)[79.9690]PSPLLSSSFDLDYDFQRDYYDR.M |

**References**


1. Villén, J., Gygi, S. P. The SCX/IMAC enrichment approach for global phosphorylation analysis by mass spectrometry. *Nat Protoc.* **3**, 1630-8 (2008).

2. Wojcik, R., Dada, O. O., Sadilek, M., Dovichi, N. J. Simplified capillary electrophoresis nanospray sheath-flow interface for high efficiency and sensitive peptide analysis. *Rapid Commun. Mass Spectrom*. **24**, 2554-2560 (2010).
3. Sun, L., Zhu, G., Zhang, Z., Mou, S., Dovichi, N. J. Third-Generation Electrokinetically Pumped Sheath-Flow Nanospray Interface with Improved Stability and Sensitivity for Automated Capillary Zone Electrophoresis-Mass Spectrometry Analysis of Complex Proteome Digests. J*. Proteome Res*. **14**, 2312-2321 (2015).
4. Zhu, G., Sun, L., Dovichi, N. J. Thermally-initiated free radical polymerization for reproducible production of stable linear polyacrylamide coated capillaries, and their application to proteomic analysis using capillary zone electrophoresis-mass spectrometry. *Talanta* **146**, 839-843 (2016).
5. Chen, D., Shen, X., Sun, L. Capillary zone electrophoresis-mass spectrometry with microliter-scale loading capacity, 140 min separation window and high peak capacity for bottom-up proteomics. *Analyst* **142**, 2118-2127 (2017).
6. Sun, L., et al. Ultrasensitive and Fast Bottom-up Analysis of Femtogram Amounts of Complex Proteome Digests. *Angew. Chem. Int. Ed*. **52**, 13661-13664 (2013).
7. Kou, Q., Xun, L., Liu, X. TopPIC: a software tool for top-down mass spectrometry-based proteoform identification and characterization. *Bioinformatics* **32**, 3495-3497 (2016).
8. Liu, X., et al. Deconvolution and Database Search of Complex Tandem Mass Spectra of Intact Proteins. *Mol. Cell. Proteomics* **9**, 2772– 2782 (2010).
9. Kessner, D., Chambers, M., Burke, R., Agus, D., Mallick, P. ProteoWizard: open source software for rapid proteomics tools development. *Bioinformatics* **24**, 2534– 2536 (2008).
10. Elias, J. E., Gygi, S. P. Target-decoy search strategy for increased confidence in large-scale protein identifications by mass spectrometry. *Nat. Methods* **4**, 207–214 (2007).
11. Tyanova S, Temu T, Sinitcyn P, Carlson A, Hein MY, Geiger T, Mann M, Cox J. The Perseus computational platform for comprehensive analysis of (prote)omics data. Nat Methods. 2016 Sep;13(9):731-40.
12. Ghandi, Mahmoud, et al. "Next-generation characterization of the cancer cell line encyclopedia." *Nature* 569.7757 (2019): 503-508.
13. McKenna, A.; Hanna, M.; Banks, E.; Sivachenko, A.; Cibulskis, K.; Kernytsky, A.; Garimella, K.; Altshuler, D.; Gabriel, S.; Daly, M.; DePristo, M. A., The Genome Analysis Toolkit: a MapReduce framework for analyzing next-generation DNA sequencing data. *Genome research* **2010,** *20* (9), 1297-1303.
14. Wang, K.; Li, M.; Hakonarson, H., ANNOVAR: functional annotation of genetic variants from high-throughput sequencing data. *Nucleic acids research* 2010, 38 (16), e164-e164.

15. Harrow, J.; Frankish, A.; Gonzalez, J. M.; Tapanari, E.; Diekhans, M.; Kokocinski, F.; Aken, B. L.; Barrell, D.; Zadissa, A.; Searle, S., GENCODE: the reference human genome annotation for The ENCODE Project. *Genome research* 2012, 22 (9), 1760-1774.
16. Chen, Wenrong, and Xiaowen Liu. "Proteoform Identification by Combining RNA-Seq and Top-down Mass Spectrometry." *Journal of Proteome Research* 20.1 (2020): 261-269.
